# Truncated Caveolin-3 Mutation In Familial Barrett’s Esophagus

**DOI:** 10.1101/2021.05.15.444305

**Authors:** Katherine S. Garman, Richard von Furstenberg, Ryan Fecteau, Thomas C. Becker, Biswa P. D. Purkayastha, Gary W. Falk, Dawn Dawson, Joseph E. Willis, Shannon J. McCall, Andrew E. Blum, Kishore Guda, Amitabh Chak

## Abstract

**Objective:** Barrett’s esophagus and esophageal adenocarcinoma demonstrate familial aggregation. The goal was to identify a segregating genetic variant in an large family and subsequently localize esophageal gene expression.

**Methods:** Whole exome sequencing of genomic DNA from affected members of a large family with Barrett’s esophagus and esophageal adenocarcinoma was analyzed to identify rare coding variants in genes segregating with disease. Histopathological assessment of archived formalin fixed esophageal human and porcine tissues to localize expression of identified genes in esophagus.

**Results:** A segregating nonsense mutation in the gene Caveolin-3 (*CAV3*) was identified. Esophageal CAV3 localized to myoepithelial cells around esophageal submucosal glands. Histologic examination of a formalin fixed paraffin embedded esophagectomy specimen from an individual carrying the *CAV3* null mutation revealed submucosal glands demonstrating atypical acinar metaplasia with absence of myoepithelial cells and no CAV3+ cells.

**Conclusions:** Submucosal glands contribute to healing of injured squamous esophagus. We theorize the truncating nonsense *CAV3* mutation disrupts normal squamous healing and the organization of submucosal glands, making affected family members susceptible to the proliferation and development of metaplastic columnar Barrett’s esophagus.

## INTRODUCTION

The majority of esophageal adenocarcinomas (EACs) originate in Barrett’s esophagus (BE), a pre-malignant metaplastic columnar epithelium, which replaces the normal stratified squamous epithelium of the injured esophagus ^1^. BE is believed to develop as a reparative response of the distal esophagus to injury from gastroesophageal reflux disease (GERD) ^1-3^. While the source of the BE cell of origin is still debated, it is widely accepted that EAC arises when BE progresses from metaplasia to dysplasia to cancer. BE and EAC are complex diseases caused by a combination of genetic and environmental factors.

BE and EAC aggregate in a proportion of families ^4^; Familial Barrett’s Esophagus (FBE), is present in 7% of probands with BE or EAC ^5^. Segregation analysis suggests that FBE is consistent with dominant transmission of one or more incompletely penetrant major Mendelian alleles ^6^.

The aim of this study was to identify a germline genetic mutation segregating with disease in an exceptionally large FBE family and to determine the cellular location where *CAV3* is expressed within the esophagus.

## METHODS

### Family Ascertainment

The NN-0001 family in this study was identified and recruited in institutional review board (IRB) approved studies at University Hospitals of Cleveland Medical Center (UHCMC) and the Hospital of University of Pennsylvania (HUP) using previously described approaches ^7^. BE cases were defined by documented intestinal metaplasia on biopsy report plus ≥ 1 cm segment of salmon-colored mucosa on EGD. EAC cases were defined as EAC on the pathology report as involving the tubular esophagus.

### Whole-exome and Sanger sequencing

Whole exome capture, library preparation, and deep sequencing were performed on germline DNA from affected living NN-0001 family members (Fig. 1) as previously described.^7^ Deep sequencing of the exome-enriched DNA pools was performed on an Illumina HiSeq 2000 instrument with 100 bp, paired-end reads to an average read-depth of 70X per sample. DNA was extracted from normal tissues contained within archived FFPE specimens from deceased NN-0001 family members and used for Sanger sequencing to confirm the presence of mutations identified through whole exome sequencing.

**Figure 1:**
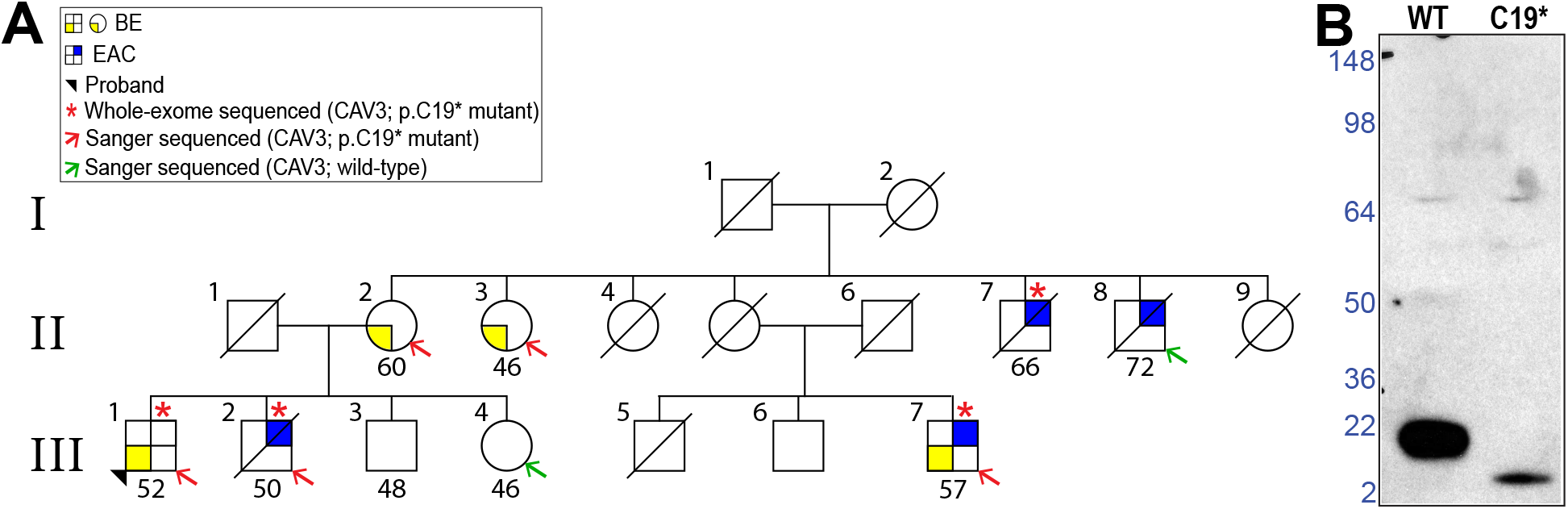
Panel A: Pedigree NN-0001 - Yellow square in left lower quadrant = BE; Blue right upper quadrant = EAC. Proband is indicated with arrowhead. Age of diagnosis, affected individuals whose exomes were sequenced, and individuals carrying the null *CAV3* mutation are indicated. Panel B: Expression of C19X null variant in mammalian cells shows greatly attenuated expression of small peptide fragment in left lane compared to wild type *CAV3* in left lane.

#### Assessing variant

To characterize sequencing variant, pcDNA4.1 expression plasmids with C-terminal FLAG-tag, containing full-length wild-type *CAV3* or *CAV3* carrying the C19* stop-gain mutant transcript, were transiently transfected into Cos7 cells, plated in DMEM media supplemented with 10% FBS. Forty-eight hours post-transfection, cells were lysed in RIPA buffer, and 40µg of total protein were resolved on 4 to 12% gel, followed by Western Blot analyses using anti-FLAG HRP antibody (1:1000), incubated overnight.

### IHC

Archival formalin-fixed paraffin-embedded tissue 5 um serial sections from human esophageal resections with normal squamous esophagus and BE/EAC were deparaffinized, rehydrated and immunostained with anti-human antibodies to caveolin 3 (CAV3), cytokeratin 5 (CK5, a marker for squamous epithelium), cytokeratin 7 (CK7, a marker for BE and columnar epithelium) and p63 (a nuclear marker for squamous differentiation and myoepithelial cells). Vendor sources, antibody dilutions and antigen retrieval methods used are designated in Table 1. Sections were incubated for 60 minutes at room temperature followed by an HRP-polymer detection system (Biocare Medical ®) appropriate for each host species was applied for 30 minutes, followed by visualization with DAB or fast RED chromagen and counterstained briefly with Gill’s Hematoxylin. Positive tissue controls (per manufacturer recommendation) and negative antibody control (antibody omission) were included.

**Table 1.** Antibodies used for Immunohistochemistry (human) and Immunofluorescence (pig)

| Target antigen | Antibody source/ cat# | Antibody type | Dilution | AR |
| --- | --- | --- | --- | --- |
| CAV3 (human) | Abcam/ ab182759 | Rabbit monoclonal | 1:100 | TRIS-HCL |
| CAV3 (pig) | Abcam/ ab2912 | Rabbit polyclonal | 1:40 | N/A |
| CAV1 (pig) | Novus/NB100-615 | Mouse monoclonal | 1:200 | N/A |
| CK5 (human) | BioCare/ CME430 | Rabbit monoclonal | 1:400 | Dako TR |
| CK7 (human) | Dako/ OV-TL | Mouse monoclonal | 1:1600 | BC* Diva |
| P63 (human) | Biocare/ CM163 | Mouse monoclonal | 1:400 | BC* Reveal |

### Porcine ESMG

Briefly, Yorkshire pigs (Sus scrofa) under North Carolina State University and Duke University IACUC146 approved protocols (NCSU 13-116-B, Duke A120-14-05). Esophagus were injured using endoscopic radiofrequency ablation (RFA) as previously described.^8^ Frozen tissue blocks were created from uninjured esophagus (City Packing, Burlington NC) and 10 days after RFA. Slides from tissue blocks were put into a solution containing 50% methanol and 50% acetone for 10 minutes at -20°C, then into 1X PBS (pH 7.4) solution for 10 minutes at room temperature, blocked with 1% BSA in PBS for 30 minutes at room temperature and incubated overnight at 4°C with primary antibodies for CAV3 and CAV1 immunofluorescence prepared in 0.1% BSA in PBS with the dilutions (Table 1). The following morning, after washing 1X PBS for 20 minutes, respective secondary antibodies prepared in 0.1% BSA in PBS were added to the slides for 45 minutes at room temperature followed by a wash and DAPI stain (1:5000) for 10 minutes. After a PBS wash, 2 drops of ProLong Gold antifade reagent and a coverslip was added. Slides were imaged and processed using the EVOS microscope and ImageJ software.

## RESULTS

### Family NN-0001 Pedigree

The proband, III-1, (**Figure 1**, arrowhead) was diagnosed with BE and high-grade dysplasia during upper endoscopy at age 52. The family reported Eastern European ancestry. The proband’s brother, III-2, had died of EAC at age 50 and mother, II-2, had BE diagnosed at age 60 and breast cancer at age 68. The mother provided the family history shown in Figure 1. Two maternal uncles, II-7 and II-8, were deceased from EAC at ages 67 and 72, respectively; a maternal male cousin, III-7, was diagnosed with BE at age 40 and underwent esophagectomy for high grade dysplasia with a T1N0 EAC at age 57; and a maternal aunt, II-3, had BE diagnosed at age 46 and breast cancer at age 57. Diagnoses were confirmed by medical record review and histological review of archived formalin fixed paraffin embedded (FFPE) biopsies from the deceased members with EAC. There was no known obesity or smoking history in the family. Family members denied any history of muscular dystrophy or cardiomyopathy, two autosomally inherited diseases associated with missense mutations in *CAV3*.^9-11^

### Identification of Segregating *CAV3* Variant

Whole exome sequencing of 4 affected individuals – III-1, III-2, III-7, and II-7 (**Fig. 1**, see asterisks) – from family NN-0001 revealed rare, i.e., reported allele frequencies < 1% in Thousand Genome, missense/splice site/indel/null variants shared by all four affecteds in 3 coding genes – *CAV3, MYO1E*, and *RCL1*. Out of these 3, the C19X null variant in *CAV3* was the only private loss of function variant, i.e. not reported in dbSNP, Thousand Genome, and allele frequency = 0.000005 in Genome Aggregation Database (gnomAD), that segregated in all four affected family members. Sanger sequencing of the shared rare and private variants found in these 3 coding genes was performed in five affected and three unaffected members of the family. The null variant in *CAV3* was the only variant that segregated in all but one of the 8 phenotyped family members. The maternal uncle who did not carry the *CAV3* null variant was the oldest affected individual in the family. For this single family, assuming a dominant one-locus two allele model at zero recombination fraction, the LOD score for the putative causative *CAV3* null variant is 0.56 (p=0.023).

### Localization of *CAV3*

Interrogation of RNA sequencing data^12^ from 18 non-dysplastic BE, 56 pre-treatment EACs, 20 normal esophageal squamous, and 11 normal gastric endoscopic biopsy samples revealed that *CAV3* was not expressed in mucosal tissues from normal esophagus, normal stomach, BE, or EAC. We also evaluated archived FFPE esophagectomy tissues from subjects with BE and EAC, and found no CAV3 in normal squamous epithelium, BE, or EAC by IHC (**Fig. 2**). We found CAV3 staining in occasional rare cells in glands at the gastroesophageal junction and at the base of multi-layered epithelium (MLE) associated with ESMGs (**Fig. 2B and 2C**).

**Figure 2:**
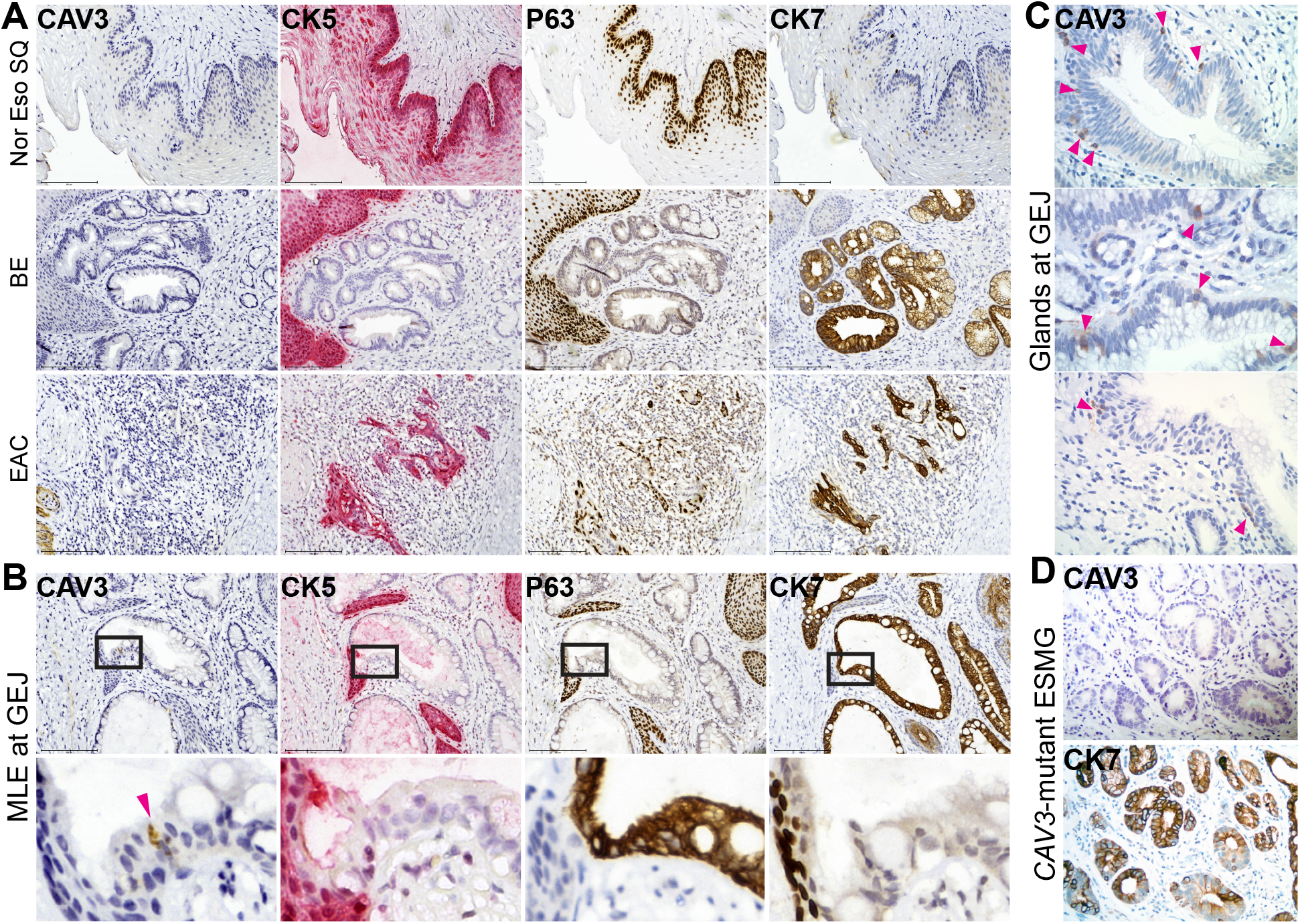
Panel A: Immunohistochemistry of normal squamous, BE, and EAC demonstrating absence of *CAV3* in all three esophageal histologies as shown in the far left column. As expected, normal esophageal squamous cells in the top row show nuclear p63 and diffuse CK5 staining. In the middle row, glandular BE (middle of each image) adjacent to squamous mucosa (far left of each image) shows CK7 immunostaining with absence of CK5 and nuclear p63. In the bottom row, poorly differentiated EAC cells show only rare areas of p63, CK5, and CK7. Panel B: Rare CAV3 positive cells, however, were noted in basal cells of transitional multi-layered epithelium (MLE). MLE region showing CK5, CK7, and p63 staining cells in the expected pattern. CAV3 positive cells appear near the transition between CK5+, P63+ squamous cells and the CK7+ columnar cells. Panel C: Rare CAV3 positive cells also noted in glands at the gastroesophageal junction and MLE associated with ESMG. Panel D: An ESMG found in an esophagectomy sample from the affected family member with *CAV3* mutant with BE and EAC demonstrates lack of CAV3 and strong patchy CK7, suggesting acinar ductal metaplasia within this ESMG.

### Histology of ESMG in Family Member with *CAV3* Variant

An archived FFPE specimen from esophagectomy performed on individual III-7 (**Fig. 2D**) who carries the C19X CAV3 variant was available for analysis. Histological examination demonstrated a striking finding of ESMGs with metaplastic acini. IHC for CK7 exhibited atypcial acinar ductal metaplasia with CK7, although CK7 was absent in the more markedly atypical acini. Notably, IHC did not find CAV3 protein in the atypical ESMGs.

### Expression of *CAV3* and *CAV1* in Normal Porcine ESMGs at Baseline and Following Injury

Typical research animals such as mice and rats lack ESMGs and it is not feasible to endoscopically collect normal ESMGs from humans. Porcine esophagus is known to contain ESMGs, so we assessed for evidence of the caveolins, *CAV3* and *CAV1* in normal porcine esophagus using immunofluorescence. In the quiescent state, CAV3 was present in selected cells in ESMGs in a distribution consistent with myoepithelial cells and CAV1 was absent (Figure 3). There were no CAV3 or CAV1 positive cells in interlobular ducts or in squamous epithelium. Ten days following esophageal RFA-induced injury, CAV3 and CAV1 were noted in dilated acini of ESMGs near the healing wound (**Fig. 3**); *CAV3* was also present throughout the healing neo-squamous epithelium.

**Figure 3:**
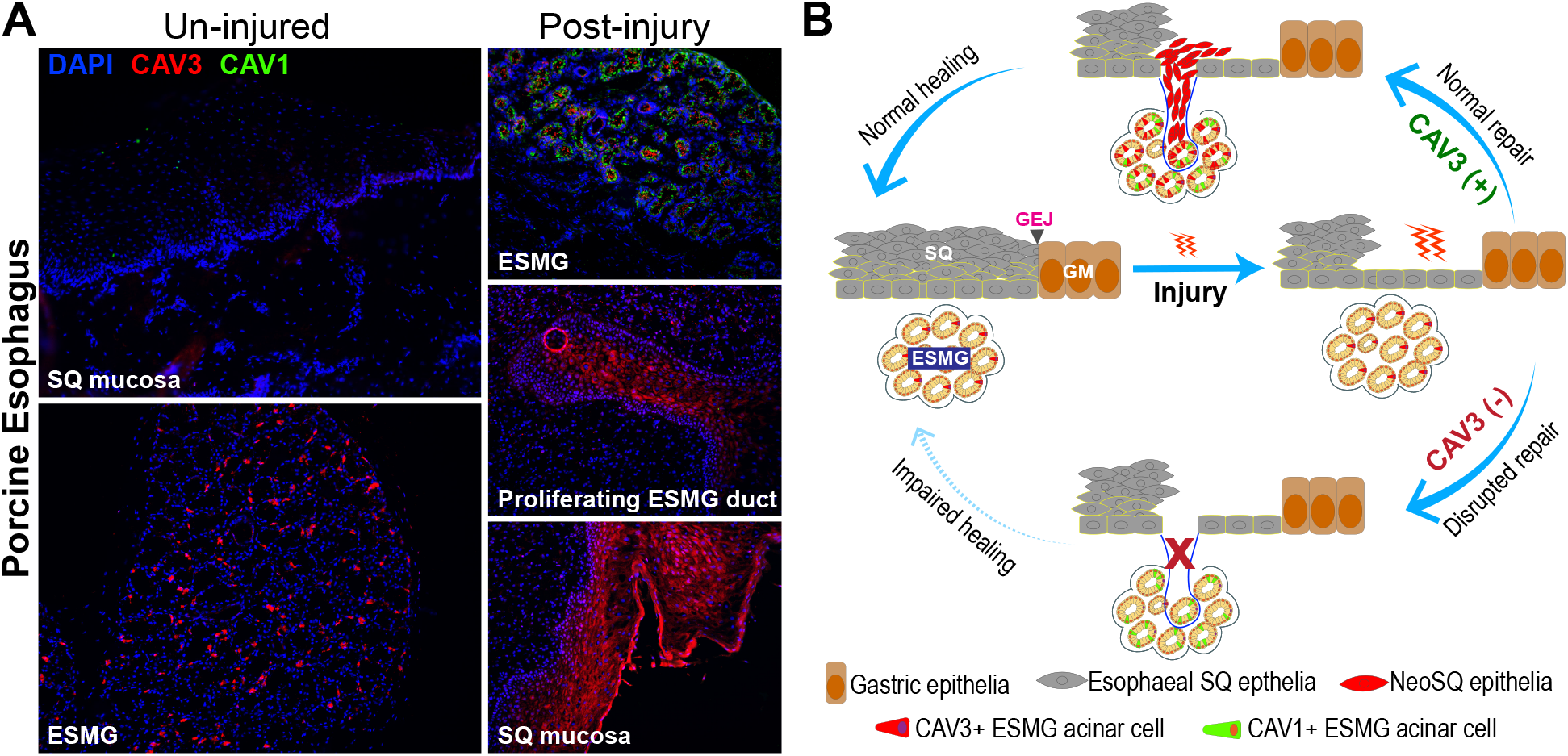
Panel A: Porcine esophagus was evaluated to detect CAV3 by immunofluorescence in uninjured esophagus. 10x images are shown with nuclear staining with DAPI in blue and CAV3 is red. Normal squamous epithelium at left did not demonstrate CAV3. In contrast, rare cells within ESMGs expressed CAV3 in the uninjured pig esophagus. CAV1 was assessed in green and was not present. Panel B: During esophageal wound healing 10 days after radiofrequency ablation, CAV3 was identified in neosquamous epithelium (left) and both CAV1 and CAV3 were identified in ESMG under the wound. Images are shown here at 10x magnification

## DISCUSSION

Whole exome sequencing of an exceptional FBE family, NN-0001, with multiple members affected with EAC and BE identified an inactivating null mutation in *CAV3* segregating with disease. Histologic examination of archived non-familial esophageal tissues plus RNA sequencing showed that *CAV3* was not expressed in squamous or metaplastic Barrett’s epithelium. We found rare *CAV3* positive cells in different niches – gastroesophageal junction glands, ESMG ducts, and base of MLE (**Fig. 2**) -- that are proposed to harbor BE precursor cells.^13^ Since *CAV3* is also known to be present in myoepithelial cells associated with glandular epithelium^14^ we examined and indeed found the presence of *CAV3* in ESMG (**Fig. 3**). Further studies of the caveolins, *CAV3* as well as *CAV1*, in the porcine injury model demonstrated increased expression of both caveolins in myoepithelial and acinar epithelial cells within ESMGs, suggesting involvement in normal squamous healing following injury. Finally, we examined an esophagectomy specimen from an affected NN-0001 family member carrying the mutated *CAV3* allele, which demonstrated a remarkably disordered ESMG. This focused our attention on the putative role of ESMGs in healing injured esophageal epithelium. We postulate that in this FBE family the loss of CAV3 function disrupts normal squamous homeostasis, which is connected to ESMGs, permitting the development BE. Genetic discovery of a loss of CAV3 function in this FBE family led to insights on the involvement of caveolins and ESMGs in normal squamous homeostasis after injury, which has just recently been demonstrated in human subjects.^15^

The caveolin, *CAV3*, is predominantly expressed in muscle and missense mutations have been associated with muscular dystrophies and cardiomyopathies.^11^ These missense mutations may cause disease by affecting oligomerization and scaffolding of *CAV3*. Family NN-0001 was found to have a protein truncating null variant and yet this family has no known muscular dystrophy or cardiomyopathy, suggesting that *CAV3* is pleiotropic.

Within the uninjured porcine esophagus, normal ESMG CAV3 localized to myoepithelial cells around acini and these cells are of particular interest given their role as tumor suppressor cells in breast as well as recent work describing the role of myoepithelial cells surrounding airway submucosal glands as reserve stem cells activated by injury^16-18^. In mice, complete loss of CAV3 has been reported to be associated with hyperplasia of mammary epithelia and increased c-Myc expression.^14^

Family NN-001 includes affected individuals with EAC, BE, and women with both BE and breast cancer, raising questions about potential myoepithelial loss and dysfunction of myoepithelial cells in this family. Esophageal histology revealed acinar ductal metaplasia within ESMGs, emphasizing the previously described associated between acinar ductal metaplasia in ESMGs with esophageal injury (ulcer), BE and EAC.^19^

*In vivo* studies have revealed that as part of the repair response following injury, ESMGs that undergo acinar ductal metaplasia have increased expression of columnar BE and progenitor markers.^20^ We hypothesize that after injury, CAV3 expressing cells may represent induced myoepithelial cells that migrate up the duct to heal the injured epithelium, similar to submucosal glands of the trachea where myoepithelial cells demonstrate the plasticity to migrate, differentiate and repair damaged epithelium ^18,21^.

We report a unique family with multiple members affected with BE/EAC. The abnormal ESMGs found in this family suggest that the CAV3 truncation in this family predisposes to BE and EAC. Further studies are needed to delineate the molecular changes at transitional and junctional epithelium or within ESMG-associated cells and determine whether alterations in the healing process after injury contributes to the development of BE and EAC rather than squamous epithelium.

## REFERENCES

1. Sharma P, McQuaid K, Dent J, et al. A critical review of the diagnosis and management of Barrett’s esophagus: the AGA Chicago Workshop. Gastroenterology 2004;127:310–30.

2. Shaheen N, Ransohoff DF. Gastroesophageal reflux, barrett esophagus, and esophageal cancer: scientific review. Jama 2002;287:1972–81.

3. Wang KK, Sampliner RE. Updated guidelines 2008 for the diagnosis, surveillance and therapy of Barrett’s esophagus. Am J Gastroenterol 2008;103:788–97.

4. Chak A, Lee T, Kinnard MF, et al. Familial aggregation of Barrett’s oesophagus, oesophageal adenocarcinoma, and oesophagogastric junctional adenocarcinoma in Caucasian adults. Gut 2002;51:323–8.

5. Chak A, Ochs-Balcom H, Falk G, et al. Familiality in Barrett’s esophagus, adenocarcinoma of the esophagus, and adenocarcinoma of the gastroesophageal junction. Cancer Epidemiol Biomarkers Prev 2006;15:1668–73.

6. Sun X, Elston R, Barnholtz-Sloan J, et al. A Segregation Analysis of Barrett’s Esophagus and Associated Adenocarcinomas. Cancer Epidemiol Biomarkers Prev.

7. Fecteau RE, Kong J, Kresak A, et al. Association Between Germline Mutation in VSIG10L and Familial Barrett Neoplasia. JAMA Oncol 2016.

8. Kruger L, Gonzalez LM, Pridgen TA, et al. Ductular and proliferative response of esophageal submucosal glands in a porcine model of esophageal injury and repair. Am J Physiol Gastrointest Liver Physiol 2017;313:G180–G91.

9. Cohen AW, Hnasko R, Schubert W, Lisanti MP. Role of caveolae and caveolins in health and disease. Physiol Rev 2004;84:1341–79.

10. Gazzerro E, Sotgia F, Bruno C, Lisanti MP, Minetti C. Caveolinopathies: from the biology of caveolin-3 to human diseases. Eur J Hum Genet 2010;18:137–45.

11. Parton RG, del Pozo MA. Caveolae as plasma membrane sensors, protectors and organizers. Nat Rev Mol Cell Biol 2013;14:98–112.

12. Blum AE, Venkitachalam S, Ravillah D, et al. Systems Biology Analyses Show Hyperactivation of Transforming Growth Factor-beta and JNK Signaling Pathways in Esophageal Cancer. Gastroenterology 2019;156:1761–74.

13. Que J, Garman KS, Souza RF, Spechler SJ. Pathogenesis and Cells of Origin of Barrett’s Esophagus. Gastroenterology 2019;157:349–64 e1.

14. Sotgia F, Casimiro MC, Bonuccelli G, et al. Loss of caveolin-3 induces a lactogenic microenvironment that is protective against mammary tumor formation. Am J Pathol 2009;174:613–29.

15. Konda V, Souza RF, Dunbar KB, et al. An Endoscopic and Histologic Study on Healing of Radiofrequency Ablation Wounds in Patients With Barrett’s Esophagus. Am J Gastroenterol 2022;117:1583–92.

16. Pandey PR, Saidou J, Watabe K. Role of myoepithelial cells in breast tumor progression. Front Biosci (Landmark Ed) 2010;15:226–36.

17. Gudjonsson T, Adriance MC, Sternlicht MD, Petersen OW, Bissell MJ. Myoepithelial cells: their origin and function in breast morphogenesis and neoplasia. J Mammary Gland Biol Neoplasia 2005;10:261–72.

18. Tata A, Kobayashi Y, Chow RD, et al. Myoepithelial Cells of Submucosal Glands Can Function as Reserve Stem Cells to Regenerate Airways after Injury. Cell Stem Cell 2018;22:668–83 e6.

19. Garman KS, Kruger L, Thomas S, et al. Ductal metaplasia in oesophageal submucosal glands is associated with inflammation and oesophageal adenocarcinoma. Histopathology 2015.

20. Kruger L, Gonzalez LM, Pridgen TA, et al. Ductular and proliferative response of esophageal submucosal glands in a porcine model of esophageal injury and repair. Am J Physiol Gastrointest Liver Physiol 2017:ajpgi 00036 2017.

21. Lynch TJ, Anderson PJ, Rotti PG, et al. Submucosal Gland Myoepithelial Cells Are Reserve Stem Cells That Can Regenerate Mouse Tracheal Epithelium. Cell Stem Cell 2018;22:653–67 e5.

